# Landscape dynamics promoted the evolution of mega-diversity in South American freshwater fishes

**DOI:** 10.1101/2021.12.13.472133

**Authors:** Fernanda A. S. Cassemiro, James S. Albert, Alexandre Antonelli, André Menegotto, Rafael O. Wüest, Marco Túlio P. Coelho, Dayani Bailly, Valéria F. B. da Silva, Augusto Frota, Weferson J. da Graça, Reginaldo Ré, Telton Ramos, Anielly Galego de Oliveira, Murilo S. Dias, Robert K. Colwell, Thiago F. Rangel, Catherine H. Graham

## Abstract

Landscape dynamics and river network rearrangements are widely thought to shape the diversity of Neotropical freshwater fishes, the most species-rich continental vertebrate fauna on Earth. Yet the effects of hydrogeographic changes on fish dispersal and diversification remain poorly understood. Here we integrate an unprecedented occurrence dataset of 4,967 South American freshwater fish species with a species-dense phylogeny to track the evolutionary processes associated with hydrogeographic events over 100 Ma. Net lineage diversification was heterogeneous through time, across space, and among clades. Three abrupt shifts in diversification rates occurred during the Paleogene (between 63 and 23 Ma) in association with major landscape evolution events, and net diversification accelerated from the Miocene to the Recent (c. 20 – 0 Ma). The Western Amazon exhibited the highest rates of *in situ* diversification and was also the most important source of species dispersing to other regions. All regional biotic interchanges were associated with documented hydrogeographic events and the formation of biogeographic corridors, including Early Miocene (c. 20 Ma) uplift of the Serra do Mar, and Late Miocene (c. 10 Ma) uplift of the Northern Andes and formation of the modern transcontinental Amazon River. Reciprocal mass dispersal of fishes between the Western and Eastern Amazon coincided with this phase of Andean uplift. The Western Amazon has the highest contemporary levels of species richness and phylogenetic endemism. Our results support the hypothesis that landscape dynamics were constrained by the history of drainage basin connections, strongly affecting the assembly and diversification of basin-wide fish faunas.

**Significance Statement:** Despite progress in mapping geographic distributions and genealogical relationships, scientists have few clear answers about the origins of South American freshwater fishes, the most diverse vertebrate fauna on Earth. Here we used the most complete dataset of geographic distributions and evolutionary relationships of South American fishes to track how the geological history of river dynamics influenced the origin, extinction, and interchange of species over the past 100 Ma. We found abrupt increases of species origination between 66 and 23 Ma, coinciding with repeated uplifts of the Andes. The Western Amazon region served as source of freshwater fishes to other regions, as a place where species tended to persist over longer historical periods, and where species originations occurred with higher frequency.

## Introduction

Geological events are widely accepted as having been central in shaping biodiversity patterns, yet we are only beginning to understand the nuanced ways in which specific historical events contributed to evolutionary diversification in most extant groups of organisms (1–4). South America harbors the most diverse freshwater fish fauna in the world (~ 5,000 species; 5,6), providing unique opportunities to study the effects of geological history and river dynamics on diversification in obligate aquatic taxa. Hydrogeographic processes operating over tens of millions of years have resulted in predictable changes in the geometry of river drainage networks, isolating and merging portions of adjacent river basins and their connections to the sea and altering the physicochemical characteristics of water discharge (7, 8). Here, we evaluate the influence of the major geological events on diversity patterns of freshwater fishes of South America over the past 100 Ma, the time period over which hydrogeographic events shaped the origins of modern fluvial systems (1, 2, 9). We generated the most complete dataset, to date, on species geographic occurrences, and in conjunction with a species-dense phylogeny of ray-finned fishes (10), we evaluated the link between hydrogeographic events and the spatial and temporal distribution of diversification and dispersal of fish clades.

The evolution of South American rivers was largely shaped by four prominent geophysical events (Fig. 1) (8, 11). The first was the final separation of South America and Africa during the Late Cretaceous (c. 100 Ma). During the Late Cretaceous and Early Paleogene (c. 100 – 55 Ma), river drainage patterns of South America were controlled by the location of the pre-existing continental uplands (cratons and shields), ongoing uplift of the Andean cordilleras, super greenhouse climatic conditions characterized by high temperatures and precipitation, and dramatically fluctuating eustatic sea levels. As a result, freshwater drainages across South America during this time were intermittently connected and isolated by changing shorelines (1, 12). During the Paleogene (c. 55 – 33 Ma), the proto-Amazon-Orinoco system (proto-Amazon hereafter) drained the Sub-Andean foreland basin, including much of northern South America and the northern La Plata region (Fig. 1a; 2, 13). Second, intraplate compression, rifting and depression of the southern Atlantic margin during the Oligocene (c. 33-23 Ma) intermittently connected and isolated headwaters of the coastal and interior drainages (Fig. 1b; 1). Oligocene plate compression also uplifted the Michicola Arch c. 30 Ma in the area of the Andean Orocline, isolating the Upper Madeira sedimentary basin from the La Plata region, and connecting it to the proto-Amazon (1). Third, uplift of Serra do Mar in southeastern Brazil during the Early Miocene (c. 23 Ma) re-routed some rivers from the La Plata basin directly to the Atlantic (Fig. 1c; 14, 15), isolating many terrestrial and aquatic species in the Atlantic Forest. Also, at about this time (c. 23-10) Ma, the Pebas Megawetland extended over large areas of the modern Western Amazon and Orinoco basins (Fig. 1c; 2, 4, 9, 16, 17). Fourth, uplift of the Northern Andes during the Late Miocene (c.10 – 4.5 Ma), which profoundly reorganized regional river drainage networks, isolated the modern Amazon, Orinoco, Magdalena, and Maracaibo basins, and connected the modern western and eastern Amazon basins, thereby forming the modern transcontinental Amazon river (Fig. 1d; 13, 18).

**Figure 1.**
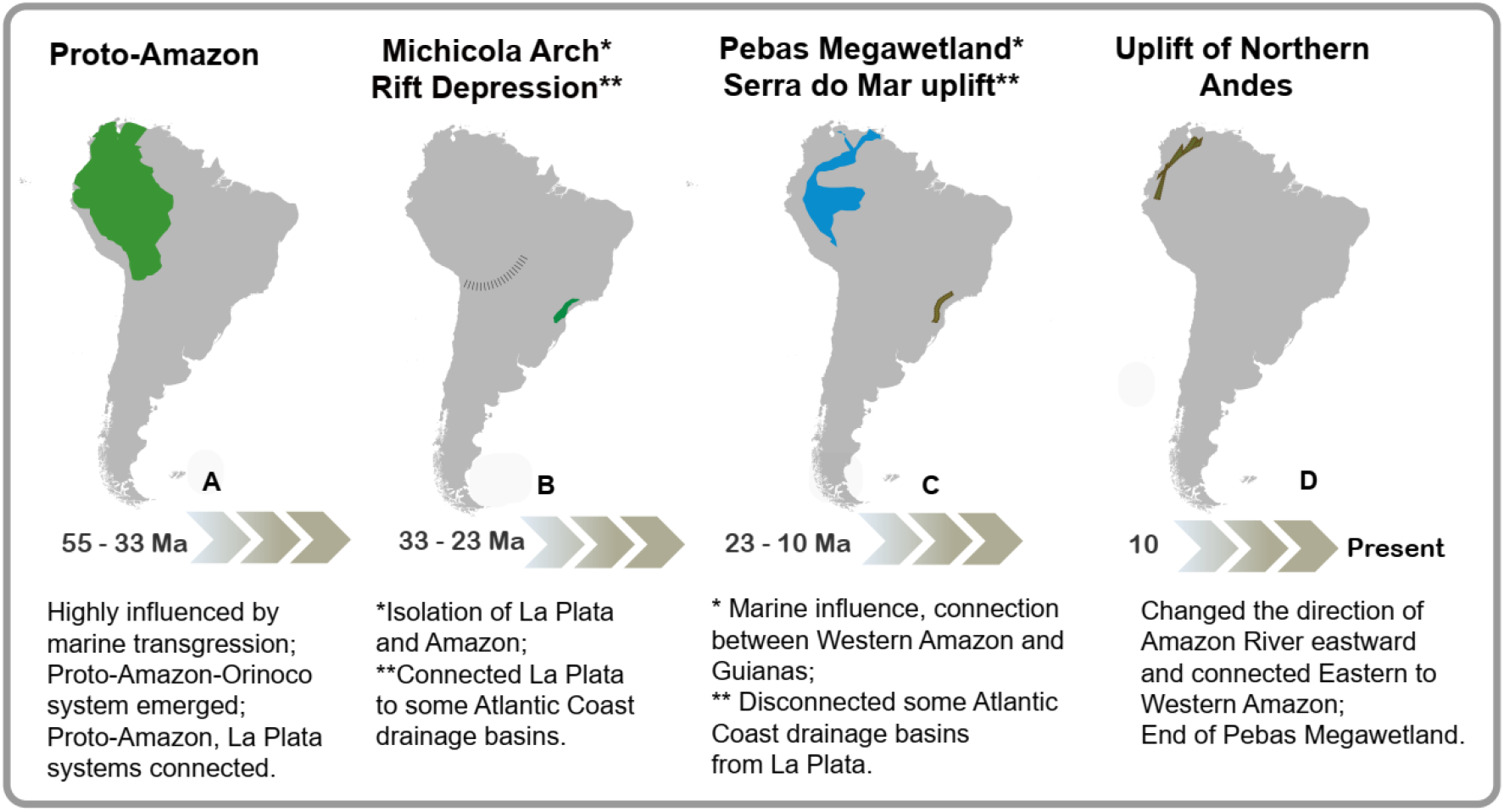
Approximate chronology and location of the principal landscape evolution events that shaped the current drainage basins of South America and influenced the diversification of freshwater fishes. Between 100 – 55 Ma, aquatic systems in South America were intermittently connected by multiple marine transgressions and regressions, thus drainages across the continent during this time were intermittently connected by epicontinental seaways. During this time, the proto-Orinoco Amazon river was the main drainage of northern South America, flowing through the sub-Andean foreland. Also, the Paraná and Paraguay basins (La Plata region; see Fig. 3) represented major aquatic systems in South America in these times (Lundberg et al. 1998, Hoorn et al. 2010, Albert & Reis 2011, Albert et al. 2018). See additional information about principal landforms controlling basin connectivity at each time interval and region delineation appears in *SI Appendix*, Table S1.

By altering the connections and configuration of regional river networks, these geological events changed genetic connections of freshwater-adapted organisms over millions of years, shaping diversity patterns of modern Neotropical freshwater fishes (1, 9). River capture is a landscape evolution process that exerts a potent influence on diversification in obligate freshwater organisms, because it both severs existing, and constructs new corridors of aquatic habitat among portions of adjacent drainage basins (19, 20). Because continental fishes are eco-physiologically restricted to freshwater habitats within drainage basins, watersheds represent natural dispersal barriers, as evidenced by the strong spatial concordance of geographic ranges in freshwater fish species with basin boundaries (21). By isolating and connecting populations of aquatic taxa across watershed divides, river capture exerts complex effects on the diversity of freshwater organisms, elevating extinction risk through geographic range contraction, promoting speciation by genetic isolation and vicariance, and increasing biotic homogenization by dispersal and gene flow (13, 22, 23).

Although geological events have long been recognized to shape the evolution of rivers and freshwater diversity, the relative contributions of particular geological events and settings remain poorly understood. Insights can be gained only by studying diversity patterns at appropriate spatial, temporal and taxonomic scales (20, 24–27). For instance, a recent study identified the Western Amazon as the center of Amazon fish diversity, with younger fish lineages dispersing progressively eastward across the Amazon after the formation of the modern transcontinental river c. 10 Ma (28). However, this interpretation overlooks the more ancient history of Neotropical fishes on the upland Brazilian and Guianas Shields, the formation of the modern lowland fauna in the proto-Amazon Sub-Andean foreland basin, and the phenotypically and taxonomically modern composition of all the known Miocene paleo-ichthyofaunas (1, 29). Taking this deeper history into account, Pliocene and Pleistocene events may be seen to have served more as buffers against extinction than as drivers of speciation in the formation of Amazonian fish species diversity (Fig. 2; 12, 30). In fact, the most species-rich clades of Neotropical freshwater fishes are thought to have radiated during the Paleogene (c. 63 – 23 Ma) (1, 31, 32).

**Figure 2.**
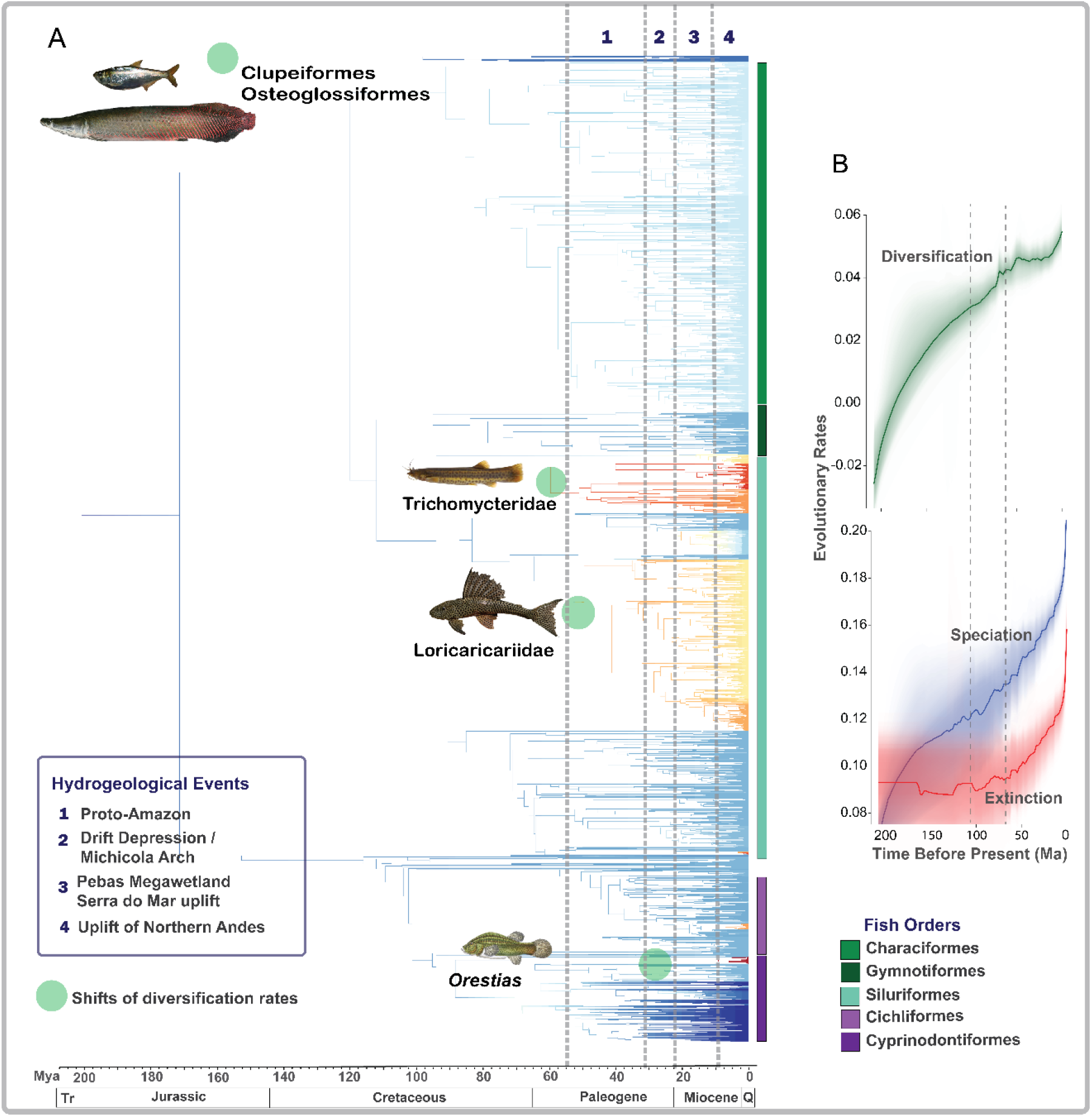
Changes in the rates of net lineage diversification among South American freshwater fishes. Tips represent 4,967 fish species. *(A)* Branch colors indicate net lineage diversification rate estimated by BAMM, where red indicates highest and blue lowest diversification rates. Significant shifts in diversification rates are shown as pale green circles on the branches. Selected representatives of orders Osteoglossiformes and Clupeiformes, families Trichomycteridae and Loricariidae, and genus *Orestias* are illustrated. The principal orders are represented by colored columns to the right of the tree tips. The timescale at the bottom is expressed in millions of years ago (Ma). Vertical dashed lines indicate timing of the main principal hydrogeographic events indicated by legend at left. *(B)* Rates-through-time plots based on BAMM estimations (see *Material and Methods* for parametrization details). The shaded areas around the curves correspond to 95% confidence intervals of the estimated rates. Dashed lines indicate the time period when most of shifts in diversification rates were estimated. Photographs: Augusto Frota, Don Stewart, and Peter van der Sleen.

In the case of South American freshwater fishes, macroevolutionary studies are hindered by the large number of species (~5000), remote sampling localities, and logistical difficulties of gathering reliable data (5). Our new data on fish distributions, which we combine with a time-calibrated molecular phylogeny, provides a unique set of resources to study the causes and consequences of past river rearrangements in shaping fish diversity, and how these geomorphological events affected diversification over longer time periods and larger spatial scales than has previously been attempted. In particular, we evaluate the prediction that the high diversity in Western Amazon was influenced by biogeographical bridges formed across different aquatic systems and time periods, which led to both accelerated diversification rates, and a role for the Western Amazon as a principal source of freshwater fish species for all of South America.

## Results and Discussion

### Diversity patterns are associated with landscape evolution

We assessed associations between landscape evolution events (e.g., tectonics, erosion, sea-level changes) with patterns of net lineage diversification (speciation minus extinction) and biogeographic dispersal (i.e., changes in species’ geographical ranges driven by landscape evolution). We used a recent species-dense molecular phylogeny of all bony fishes (10, 33) to estimate net lineage diversification rates and rate shifts through time (34). We estimated phylogenetic endemism of each drainage basin (35) using a new and unprecedentedly complete dataset comprising ~95% all fish species described for South America. Basins were defined using a high-resolution SRTM digital elevation model (36). We conducted ancestral area estimation of freshwater fishes using the dispersal-extinction-cladogenesis (DEC) model of geographic range evolution in the software platform BioGeoBEARS (37, 38) to evaluate fish dispersal events among six previously defined regions, based on taxonomic similarity (see *SI Appendix*, Biogeographical Regionalization section and Fig. S1). We interpret these biogeographical reconstructions, taking into account known events of expansion, contraction, and connectivity of continental drainage basins. As fish dispersal is limited by drainage basin boundaries, it is important to reconstruct dispersal events to understand how and when diversification processes took place.

Our results show that high numbers of species and endemic species are strongly associated with processes of landscape evolution, which both constrained and promoted species interchange across basins over millions of years. Considering all dispersal events (17,244) from all regions, most (9.8%) occurred from the Western Amazon to the La Plata regions (Fig. 3; *SI Appendix*, Table S2, S3), facilitated by intermittent connections across the low-elevation watershed of the Upper Madeira and Upper Paraguay basins. Subsequently the Late Miocene uplift of the Northern Andes breached the Purus Arch, facilitating the connection of the Western and Eastern Amazon and increasing species interchange between these two large, low-elevation and species-rich regions. In addition, some basins along the Atlantic Coast have been isolated for the last 25 Ma, contributing to high fish endemism of this region. Biogeographical corridors between Western Amazon and other regions were fundamental in the build-up of fish diversity across South America.

**Figure 3.**
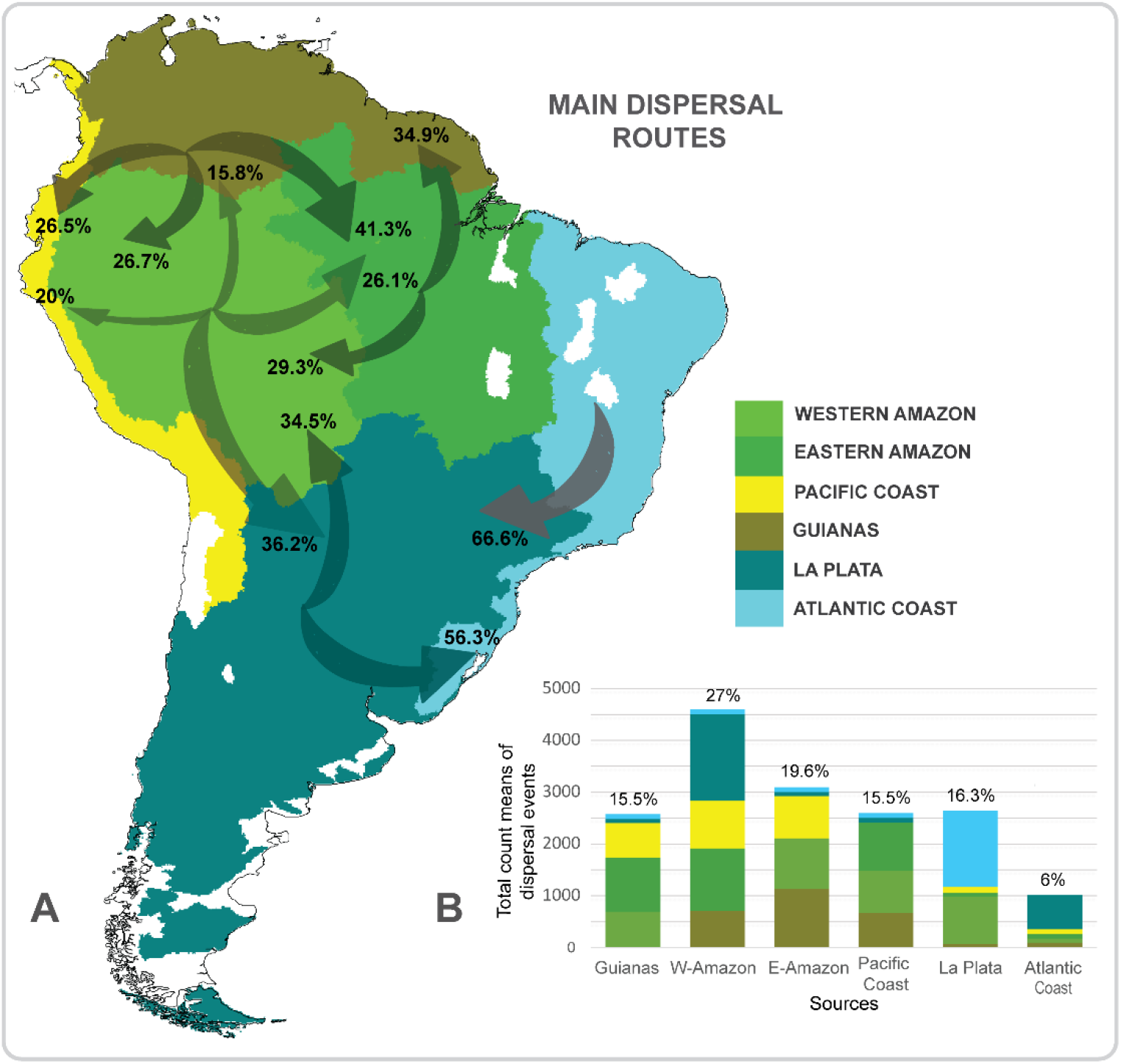
*(A)* Proportions of biogeographical dispersal events among regions. Arrows indicate dispersal directions, and numbers represent percentage of the mean (of total 20 simulations) for all dispersal events from source to sink regions. Width of arrows is proportional to the number of dispersal events. *(B)* Total count means of dispersal events from source (axis x) to destination regions (colored legend). Percentages over each column represent the relative contribution of each source region to total number of dispersal events. The drainage basins in white were not considered in the study for lack of adequate sample effort. For more details regarding dispersal analysis results, see *SI Appendix*, Tables S2, S3.

### Diversification patterns

We found that the diversification of freshwater fish in South America was heterogeneous through time, across space and among taxonomic groups (Fig. 2). Diversification results based on Bayesian analysis of macroevolutionary mixtures (BAMM; 34 – see *Material and Methods* for details) for 4,967 species, revealed four rapid shifts in diversification rate, mostly between the Late Cretaceous and the Paleogene (70-30 Ma) (Fig. 2a). During this period, the region of the proto-Amazon was largely connected and highly influenced by marine transgressions and regressions, and conversely, the retraction and expansion of freshwater habitats (1, 2, 31). These dynamics are likely to have promoted suitable conditions for speciation, enhanced by the repeated influx of marine species combined with periods of isolation of freshwater habitats (5, 31, 39). Paleontological data corroborate the hypothesis that Paleogene radiations of freshwater fishes occurred in the proto-Amazon region (39). These dynamics likely caused the high contemporary level of evolutionary distinctiveness (the amount of unique evolutionary history a lineage represents), especially in basins located along the Amazon River and in the southernmost basins in the La Plata region (Fig. S2). Our results are also consistent with the Paleogene radiation hypothesis, revealing three diversification shifts at that time (Fig. 2b). This period showed shifts in diversification rates in the clades Trichomycteridae and Loricariidae, both broadly distributed across tropical South America (see 40; *SI Appendix*, Fig. S3), and *Orestias*, restricted to high-elevation habitats in the Andes (41). Amazonian genera such as *Arapaima, Osteoglossum, Anchoviella* and *Pellona* (Osteoglossiformes and Clupeiformes) experienced shifts in diversification in the Early Jurassic (c. 160 Ma) (Fig. 2a; *SI Appendix*, Fig. S3).

Rates of diversification, especially for the species-rich clades Trichomycteridae and Loricariidae, began to accelerate about 30 Ma (Fig. 2a). First, the rise of Michicola Arch (c. 30 Ma) blocked the connection between proto-Amazon and La Plata region (1). Then, the Pebas Megawetland (c. 23 - 10 Ma) joined Western Amazon and Orinoco (Guiana region) drainages (2, 12, 13). At the same time, the Purus Arch blocked dispersal between Western and Eastern Amazon, isolating these regions from c. 40-10 Ma (1, 2). Next, rapid uplift of Northern Andes (c.10 Ma) caused drastic changes in the drainage network, changing the direction of the Amazon River flow from westward to eastward, connecting the Western and Eastern Amazon. Finally, high speciation rates in the last several million years could have been influenced by the Uplift of the Serra do Mar, which isolated the La Plata basin from some Atlantic Coast drainage basins, an event correlated with an acceleration of speciation by vicariance (Fig. 2a).

### Routes and timing of dispersal

Our biogeographical analyses showed that the Western Amazon is the most important source of freshwater fish lineages that originated in the Miocene, among the five regions we considered in South America (Fig. 3a, b; *SI Appendix*, Tables S2, S3). We recorded a total of 17,244 estimated dispersal events across South America, of which the Western and Eastern Amazon together contributed almost half (46.6%) of the total. The Western Amazon contributed the most species to other regions (27% of dispersal events considering all regions; Fig. 3b), with the largest number of dispersal events toward to the La Plata region (36.2%; Fig. 3a). The Eastern Amazon region was the next most common source (19.6%; Fig. 3b), with most of these dispersal events to the Guianas region (Fig. 3a). In addition, we estimated the relative probability of dispersal among regions through time, from 210 Ma to present, based on our hypothesized principal hydrogeographic events (Fig. 1 and *SI Appendix*, Table S1 for more details). Dispersal from the Western Amazon to the La Plata region peaked twice, first around 40 Ma followed by an abrupt decrease around 30 Ma, while the second peak occurred between 10 and 5 Ma. Dispersal from Western to Eastern Amazon regions began around 10 Ma (Fig. 4).

**Figure 4.**
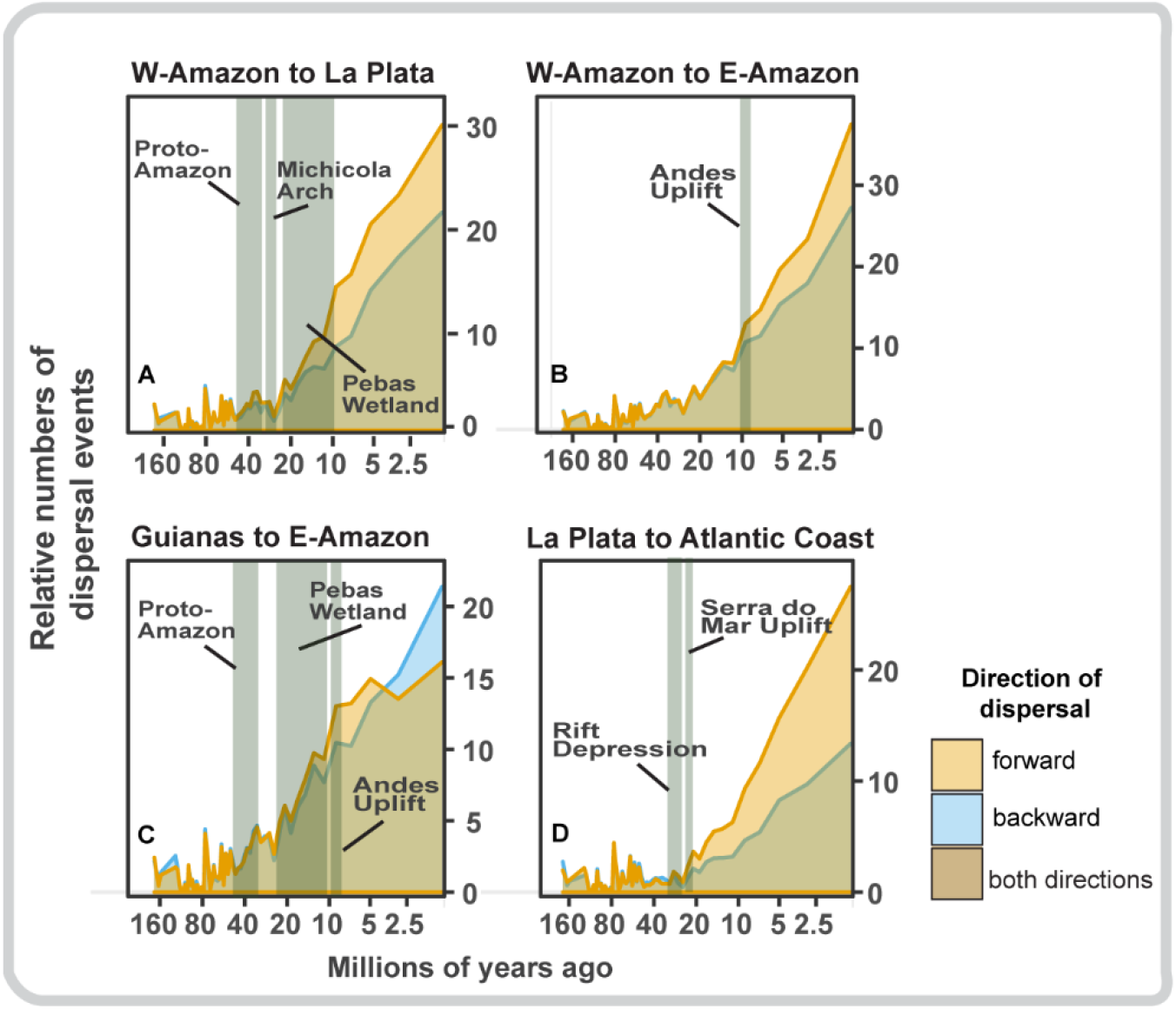
Principal dispersal events through time among the regions that acted most prominently as species sources, and geological events that influenced the connections among drainage systems, triggering the dispersal events across South America. We depicted, here, only dispersal events since 160 Ma, but older events were included in the total calculation of events. The hydrogeographic events considered follow those represented in Figure 1. The relative number of dispersal events is the total number of dispersals divided by the log-number of lineages within each time bin. The time scale on the X-axis is logarithmic.

Although the history of landscape connections between the Western Amazon and La Plata basins is not entirely clear, they likely involved multiple river capture events over millions of years (1). The fish faunal interchange around 40 Ma between these regions (Fig. 4a) may have been enhanced by mega river capture events in the sub-Andean foreland in the vicinity of the Bolivian Orocline (13, 20). The abrupt decrease in dispersal between these regions temporally coincides with the rise of the Michicola Arch (c. 30 Ma; Fig. 4a), an important dispersal barrier for riverine fishes (42). Although the beginning of the second peak of dispersal coincides with the origin of the Pebas Megawetland (c. 23 - 10 Ma; Fig. 4a), there is no geological evidence that Western Amazon and La Plata regions were connected during this time (43). Although the La Plata has not been permanently connected to the Western Amazon for at least 10 My, a semipermeable dispersal corridor between the two systems still emerges when there are intense, seasonal floods, allowing for movement of species adapted to the environments of ephemeral streams and marshes (44). This ephemeral connection explains at least some of the most recent dispersal events (45; Fig. 4a), supporting the hypothesis that the Paraguay basin (La Plata) harbors more clades from Amazon basin than the contrary (44, 46), and also provides quantitative evidence for the hypothesis that similarities in species composition between Amazon and the headwaters of the Paraguay basin (La Plata) are due to connections that predate the formation of the modern Amazon watershed (44). Other important past drainage connections are highlighted by our results, including an increase in the frequency of dispersal events between Western and Eastern Amazon (Fig. 4b). This increase coincided in time with the most recent uplift of the Northern Andes, starting c. 10 Ma, which changed the direction of Amazon River flow and connected these two regions.

The formation of the Pebas Megawetland (c. 23 Ma; 2) expanded the proto-Amazon drainage area westward, and some proto-Orinoco tributaries (Guianas) started discharging to the Eastern Amazon (1, 13), facilitating dispersal from the Guianas to the Eastern Amazon during the Miocene (Fig. 4c). The most recent uplift of the Northern Andes (c. 10 Ma) created a divide between the upper Amazonas and Orinoco drainage systems, which explains the subsequent decrease in dispersal. However, this divide is permeable, with a dispersal corridor through the Casiquiare channel during the rainy season, allowing some fish species to move from Eastern-Amazon to the Guianas (47). Previous studies have suggested that the lineages within the Orinoco and Amazon basins appear to share common ancestors more recently than 8 Ma (48, 49). Loricariid (50) and cichlid (51) species currently distributed in both regions offer potential examples of dispersal between these regions.

The peak of dispersal (in all directions) between the Atlantic Coast and La Plata regions occurred c. 70 Ma, with subsequently slightly higher dispersal events from the Atlantic Coast to La Plata (c. 60 – 40 Ma) (Fig. 4d). This pattern of dispersal corroborates the hypothesis of intense faunal interchange from upland to lowland (including continental rivers) of the Atlantic Coast drainage systems during this period, especially in the Northeastern coast, with subsequent diversification (14). Some fossils of fishes that inhabit large rivers of the La Plata basin are found in the Tremembé Formation (14), a sedimentary formation in the Ribeiro de Iguape basin of the Atlantic Coast. Although the rift depression resulted in a connection between the La Plata and some Atlantic Coast basins in southeast Brazil around 30 Ma, this event did not have a strong impact on dispersal events as a whole, but probably promoted diversification processes, resulting in higher levels of endemism (see also Fig. 5b). Subsequent to the uplift of the Serra do Mar, most Atlantic Coast basins flowed directly to the sea, with dispersal corridors between these two regions persisting in the northeast of Brazil (14). This pattern explains the high number of dispersal events from La Plata to the Atlantic Coast region (from c. 20 Ma to the present). The hypothesis of recent connections between the La Plata and Atlantic Coast regions is supported by the presence of several extant species distributed on both sides of the current watershed divide (52, 53).

**Figure 5.**
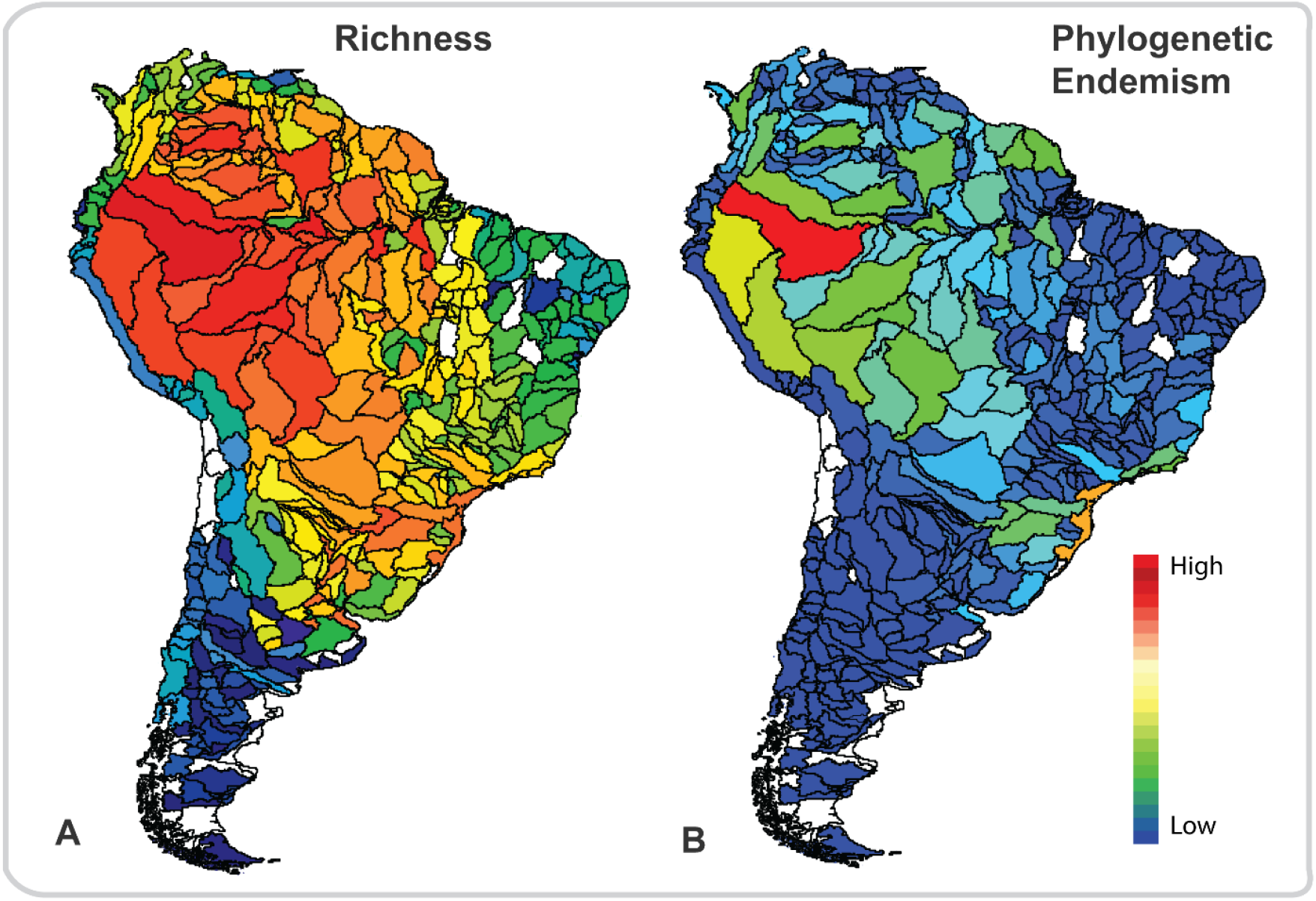
(A) Species-density map. Density was calculated as Species/(Area)^b^, where the value of *b* was calculated using the Species-Area Relationship. (B) Phylogenetic endemism (mean) for each drainage basin. Phylogenetic endemism, a combination of the phylogenetic diversity and weighted endemism measures, identifies regions to which unique phylogenetic lineages are restricted. The drainage basins in white were not considered in the study for lack of adequate sampling effort (see *Methods* for details).

### Contemporary species richness and phylogenetic endemism patterns interpreted in the light of historical events

Freshwater fish species richness (expressed as a species-density map) showed a clear latitudinal and longitudinal pattern across South America. Species richness is lower in higher latitudes and higher near the Equator (Fig. 5a). As a consequence of the effects of past hydrogeographic events on evolutionary and ecological processes, the richness and endemism of freshwater fish in the Western Amazon are extremely high (Fig. 5a, b), increasing in the opposite direction of the river flow (54). This pattern is a result of the uplift of the Northern Andes and breaching of the Purus Arch, which promoted dispersal between the Western and Eastern Amazon and isolation in headwaters region of the Orinoco (Guianas) and Western Amazon basins (Fig. 3, Fig. 4). These landscape dynamics enhanced net-diversification rates (Fig. 2) and, consequently, increased the diversity and endemism of the Western Amazonian fish fauna (3). In contrast, marine incursions in the Eastern Amazon since the Late Cretaceous likely reduced species richness near the mouth of Amazon River (13, 54). Our results differ markedly from the general pattern observed in rivers worldwide, described as the river continuum concept, in which diversity is expected to increase from headwaters to mouth (54–56).

Past biogeographical barriers can explain the high level of phylogenetic endemism in the basins of the southeastern Atlantic Coast (Fig. 5b). The isolation of some Atlantic Coast basins from La Plata basins, driven by the uplift of the Serra do Mar, provided stable relict habitats and opportunities for vicariance at higher elevations. These areas have recently been described as historically stable refugia for diverse taxa in the Atlantic Forest (57, 58) and hold high levels of genetic diversity and endemism for several groups (14, 59, 60).

### Conclusion

Advances in the biogeography of South American fishes have been hampered by poor sampling and by the concentration of most studies on small taxonomic groups, within a limited spatial scale, especially in the Western Amazon and Guianas Shield (6, 13, 61, 62). Here, we minimized the effects of poor sampling by employing a new simulation procedure to estimate species presence/absence in each basin, given sampling effort biases, following principles of existing statistical methodology that accounts for uncertainty and uneven sampling. In so doing, we were able to offer a more reliable evaluation of the relationship between landscape evolution and freshwater fish diversity.

Our study shows that shifts in diversification rates predate the Andean uplift and were not confined to the modern Amazon Basin. Rather, a combination of biogeographic and phylogenetic evidence indicates that many fish lineages of the Western Amazon are older and dispersed over much of South America (6, 62). Our results reinforce the hypothesis that landscape dynamics that played out over the last 100 My were fundamental to the establishment of drainage basins, creating both dispersal routes and barriers, thus contributing to the assembly of basin-wide faunas (8, 12, 63). Hydrogeographic events caused isolation and mixing of preexisting aquatic faunas through a series of vicariance and dispersal events, resulting in a complex history of speciation within and between basins. The Western Amazon differs from other basins in presenting high levels of phylogenetic endemism and diversity, acting as both museum and cradle of species (64), as well as source of species for other regions. In recent decades, alarming rates of deforestation, land-use degradation, damming and water pollution in South America (65, 66) have driven the potential loss of many aquatic species, especially in the Amazon and the Atlantic Forest regions, (67, 68). Knowledge of the evolutionary processes underpinning such diversity promises to help identify priority areas for conservation and focus strategies to maintain the diverse and highly endemic fish fauna of these areas.

## Materials and Methods

### Occurrence and phylogenetic data

We compiled 306,425 occurrence records of 4,967 freshwater bony fish species in South America, through an extensive search in the literature, museums, FishBase (http://www.fishbase.org/), FishNet (http://www.fishnet2.net), Global Biodiversity Information Facility (GBIF) (http://www.gbif.org/), SpeciesLink (http://splink.cria.org.br), and Datos de Peces de Aguas Continentales de Argentina (http://www.pecesargentina.com.ar/). We automated the process of data filtering by writing and implementing code that considered all possible errors in those databases (e.g., georeferencing errors, synonymies, collection year, typos, exotic species). We removed all records of species that are exotic to South America, based on Gubiani et al. (69). Species names were validated following the Catalog of Fishes of the California Academy of Sciences (http://www.calacademy.org/scientists/projects/catalog-of-fishes) and carefully revised by taxonomists specialized in Neotropical fishes. Occurrence records were then mapped into 490 drainage basins (one of the units of analysis in this study), as delineated by the HydroBASINS database, a subset of HydroSHEDS project (level 5, Hydrological data and maps based on Shuttle Elevation Derivatives; (36), available at https://hydrosheeds.org/page/hydrobasins), to estimate spatial patterns of species richness and diversification rates across South America. The HydroBASINS database provides a range of drainage basin resolutions, from level 1 (low) to 12 (high resolution). We chose level 5 as an intermediate resolution level that is adequate for the sampling effort within sub-basins.

We used the time-calibrated molecular phylogeny of actinopterygian (ray-finned) fishes constructed by Rabosky et al. (10); see also the package *fishtree*; 32, and http://fishtreeoflife.org) which includes data for all 4,967 freshwater fish species used in our analyses. As Rabosky et al. (10) proposed 100 phylogenetic tree solutions, we selected the tree with the Maximum Clade Credibility (MCC) for further analyses. As the tree by Rabosky et al. (10) presents some taxonomic incongruences, we corrected the relationships within Ostariophysi, according to Hughes et al. (70). Thus, we have the corrected position of clades as follows: (Characiformes(Gymnotiformes(Siluriformes))).

### Filling the gap in species distributions

The mapping of species records in drainage basins revealed a heterogenous sampling effort across South America, with as much as a 10-fold difference in number of occurrence records and 5-fold difference in species richness in basins of comparable area (see *SI Appendix*, Fig. S4). To mitigate sampling heterogeneity, we employed a novel simulation procedure to estimate species presence/absence given sampling effort biases, following principles of existing methodology that accounts for uncertainty. The approach consists of two iterative steps, which we repeated independently for each focal basin. We first calculated a completeness index for each drainage basin (level 5) (71, 72), by estimating the probability that the next record would add a new species to those recorded in the basin (the “coverage deficit” of 71). If the lower 95% confidence limit of the completeness index was greater than or equal to 0.75, and the number of unique records (i.e., unique combination of species name, geographic coordinates and sampling date) was larger than 50, the sampling effort at the focal drainage basin was considered acceptable. However, if the completeness index was less than 0.75, we randomly sampled an occurrence record among all adjacent drainage basins, i.e., level 5 basins in the same larger catchment delineation, level 4. We adopted this conservative criterion because random effects may turn the completeness estimate unstable in basins with small numbers of records, returning artificially high completeness values (73). Also, resampling from adjacent basins that belong the same large basin (level 4), conservatively assumes low dispersal capacity and does not increase the potential species pool. The procedure then iterates back to step one, sampling again, with replacement, after each new occurrence record is inserted into the focal basin, until the completeness index reaches at least 0.75. We repeated this procedure 100 times for all South American drainage basins to account for uncertainty, thereby generating 100 matrices of species presence/absence. We excluded from analysis 20 basins that were poorly sampled (white basin on map) and did not have an adjacent basin (i.e., additional level 5 basin within level 4 catchment region; see *SI Appendix*, Fig. S4).

### Analysis of Phylogenetic Endemism

We estimated the Phylogenetic Endemism (PE) of each drainage basin (34). PE uses the branch length and the geographic range of the extant descendants of a clade to apportion Phylogenetic Diversity (PD) across the areas where it occurs, therefore identifying concentrations of spatially restricted PD (74, 75). See *SI Appendix* for more details.

### Evolutionary rates

Diversification, speciation and extinction rates were estimated using the Bayesian analysis of macroevolutionary mixtures (BAMM; 33), considered an accurate approach among different evolutionary tests (see 76). We performed BAMM with three runs each with 1 million Markov Chain Monte Carlo (MCMC) generations and sampling frequency of 1000. We discarded a burn-in of 10% and confirmed that effective sample size values were appropriate. Results were analyzed and plotted with various functions in BAMMtools as follows. We derived prior settings from R library BAMMtools, using the function setBAMMpriors before the analysis and modified the default setting to achieve convergence. The priors used were expectedNumberOfShifts = 1.0; lambdaInitPrior = 5.32213784027579; lambdaShiftPrior = 0.00553434074897555; muInitPrior = 5.32213784027579; Prior of the time mode being time-variable (vs. time-constant). The prior parameterization ensures that the prior density on relative rate changes across the tree is invariant to the scale of the tree (76). We used computeBayesFactors to identify the best-supported model of rate shifts in our data. We plotted the maximum a posteriori probability shift configuration (getBestShiftConfiguration), the 95% credible set of distinct rate shift configurations (credibleShiftSet), and a ‘phylorate’ graph showing mean marginal posterior density of diversification rates (plot.bammdata) for each lineage. Rates-through-time plots were also generated for speciation (λ), extinction (µ), and net diversification (*r*) using ‘PlotRateThroughTime’. We are aware of potential limitations regarding BAMM, which has been criticized for estimating unrealistic extinction rates (77, 78), however, BAMM’s proponents responded that this criticism was based on spurious premises (10, 76).

Briefly, the criticism relied on the priors used by the software, but we adjusted the priors using the latest version (see BAMM documentation), in which this issue was resolved. Other problems cited by that study can be applied to most macroevolutionary methods (e.g., estimation of extinct clades) and in this sense BAMM was not considered better or worse than similar software (76).

Lastly, one of the advantages of BAMM and similar methods (79–83) is that lineages can differ in their rates of speciation and extinction, that is, the rates can vary among lineages and though time. Thus, BAMM allows us to describe multiple processes that explain rates of diversification on different parts of the tree.

### Ancestral range reconstruction and dispersal events

We reconstructed the biogeographic history of freshwater fishes, considering the delimitation of 6 regions of South America, which broadly correspond to regional drainage basins (level 2 of HydroBASINS; see details of regions delimitation analysis in *SI Appendix*). To infer geographical range evolution of lineages (i.e., ancestral range analyses; see *Appendix 1* regarding the evolution of ancestral ranges), we applied a dispersal-extinction-cladogenesis model (DEC) implemented in BioGeoBEARS (36; 82). We analyzed dispersal events by taking into account biotic connectivity among all regions. Connectivity was modeled based on paleogeographic and geological events (Fig. 1; 1, 2, 13) incorporating time-stratified dispersal multiplier matrices in the model (*SI Appendix*, Table S1). We divided the hydrogeographic history of South American freshwater fishes, from 210 Ma to present into five strata: 210–33, 33-23, 23-10, 10-5, and 5 Ma to present, according to the principal hydrogeological events mentioned in Fig 1 (see *SI Appendix*, Table S1 for more details). The dispersal multiplier matrices for each of these strata allow estimation of the relative probability of dispersal between regions for each time stratum and are roughly scaled to represent the connectivity between the areas during each time stratum. We computed the absolute number of dispersal events through time by extracting the areas and ages of all nodes from MCC phylogenetic tree, using the Biogeographical Stochastic Mapping approach (84). As the number of lineages (branches) in any phylogeny tends to increase through time, increasing the number of dispersal events toward the present even under a constant dispersal rate, we calculated the relative numbers of dispersal events by dividing absolute numbers by the total number of all lineages within each time bin. However, there are extremely few branches towards the tree root, which causes an over-representation of (relative) dispersal towards the root. To reduce the impact of that bias, the number of dispersals per time bin was divided by the log-number of lineages (per the same time bin). A full description of the methodology used, appears in *Supplementary Information, Appendix*.

## Acknowledgments

FASC is supported by CAPES postdoctoral fellowship and a visiting fellowship of Federal Institute for Forest, Snow and Landscape Research (WSL). JSA acknowledges financial support from United States National Science Foundation awards DEB 0614334, 0741450, and 1354511. AA is supported by funding from the Swedish Research Council, the Swedish Foundation for Strategic Research, and the Royal Botanic Gardens, Kew. ROW and CHG acknowledge funding from the European Research Council (ERC) under the European Union’s Horizon 2020 research and innovation programme (grant agreement No 787638). AF thanks to the National Council for Scientific and Technological Development for granting postgraduate scholarship (CNPq 141242/2018-3). WJG is grateful to the National Council for Scientific and Technological Development (CNPq/Ministry of Science, Technology, Innovations and Communications) by the research productivity grants (PQ No 305200/2018-6).

## Notes

**Competing Interest Statement:** The authors declare that they have no conflict of interest.

### Competing Interest Statement

The authors have declared no competing interest.

