## Supplemental Material for "Landscape dynamics promoted the evolution of mega-diversity in South American freshwater fishes"

**This PDF file includes:**

- Materials and Methods
- Figures S1 to S4
- Tables S1 to S3
- Appendix 1
- Supplementary References

### Materials and Methods

**Principal landforms controlling basin connectivity at each time interval.** We relied on Lundberg et al. (1998), Hoorn et al. (2010a,b), Albert & Reis (2011), Cogne et al. 2012, Tagliacollo et al. 2015, Ribeiro (2016), Jaramillo et al. 2017, Albert et al. (2018), Bicudo et al. (2019) and Bernal et al. (2019) to define the approximate chronology and location of the principal landscape evolution events that shaped the current drainage basins of South America and influenced the diversification of freshwater fishes. We defined, for each time interval, the possible connections among the six regions defined by the regionalization analysis (see below), as shown in **Table S1**.

**Table S1.** Paleogeographic model used for the time-stratified biogeographic analyses implemented in our dispersal–extinction–cladogenesis (DEC) model. The model is built on connectivity matrices among regions, with connectivity changing through time. For any combination of two regions, each matrix designates whether the areas are connected (1) or disconnected (0). When two regions are connected, dispersal events and range expansions are allowed in the biogeographic reconstruction. When two regions are connected, an ancestor can expand its range across these regions and adjacent regions connected to them. This simple model is based on current knowledge of South American hydrogeological evolution but should be viewed as a working hypothesis to be further refined and tested. (**Guianas**: Orinoco, Guianas and some trans-Andean drainages. **Western-Amazon**: From Western Amazon drainages to the area of pre-existent Purus Arch. This region currently encompasses all drainage basins of the Amazon River. **Eastern-Amazon**: From Purus Arch to the Atlantic Ocean, where currently the mouth of Amazon River is located. **Pacific Coast**: Most drainage systems that drain to the Pacific Coast. **La Plata**: Drainage systems encompassing areas currently drained by Paraná and Paraguay Rivers, besides Patagonia area. **Atlantic Coast**: Southeastern and Northeastern Atlantic drainage systems. All regions were defined according to the analysis described in section “Biogeographical Regionalization”

| **Regions** | **Guianas** | **Western-Amazon** | **Eastern-Amazon** | **Pacific Coast** | **La Plata** | **Southeastern Coast** | **Principal Landscape Events** | **References** |
|---|---|---|---|---|---|---|---|---|
| **55 - 210 Ma** | | | | | | | | |
| | Aquatic systems in South America were intermittently connected by multiple marine transgressions and regressions, thus drainages across the continent during this time were intermittently connected by epicontinental seaways | | | | | | All regions were considered connected | Lundberg et al. 1998, Lundberg et al. 2010 |
| **Eocene: 55 – 33 Ma** | | | | | | | | |
| **Guianas** | 1 | 1 | 1 | 1 | 0 | 0 | Sub-Andean Foreland: connected La Plata (Paraguay) and Western-Amazon | Tagliacollo et al., 2015, Albert et al. 2018 |
| **W-Amazon** | 1 | 1 | 1 | 1 | 1 | 0 | Proto-Amazon: connected Eastern-Amazon and La Plata | Hoorn et al., 2010a,b, Albert et al. 2018, Tagliacollo et al. 2015 |
| **E-Amazon** | 1 | 1 | 1 | 1 | 0 | 0 | Low altitude of the Northern Andes | Hoorn et al., 2010; Lundberg et al. 1998 |
| **Pacific Coast** | 1 | 1 | 1 | 1 | 0 | 0 | | |
| **La Plata** | 0 | 1 | 0 | 0 | 1 | 0 | | |
| **Atlantic Coast** | 0 | 0 | 0 | 0 | 0 | 1 | | |
| **Oligocene: 33 – 23 Ma** | | | | | | | | |
| **Guianas** | 1 | 1 | 1 | 1 | 0 | 0 | | |
| **W-Amazon** | 1 | 1 | 0 | 1 | 0 | 0 | Rise of Michicola Arch: disconnected Western-Amazon and La Plata | Tagliacollo et al., 2015 |
| **E-Amazon** | 1 | 0 | 1 | 1 | 0 | 0 | Breach Garupa Arch: when the Amazon river began flowing over the Garupa Arch, the Central Amazon (CA) became connected to the Eastern Amazon (EA). | Hoorn et al., 2010, Lundberg et al. 1998 |
| **Pacific Coast** | 1 | 1 | 1 | 1 | 0 | 0 | | |

| | | | | | | | | |
|---|---|---|---|---|---|---|---|---|
| **La Plata** | 0 | 0 | 0 | 0 | 1 | 1 | Rift Depression indicates connection between La Plata and Southeastern Coast | Ribeiro 2006, Lundberg et al. 1998 |
| **Atlantic Coast** | 0 | 0 | 0 | 0 | 1 | 1 | | |
| **Early-Middle Miocene 23 – 10 Ma** | | | | | | | Pebas Megawetland: diversification in Proto-Orinoco-Amazon | |
| **Guianas** | 1 | 1 | 1 | 1 | 0 | 0 | Pebas Megawetland extended over large areas of the modern Western Amazon and Orinoco basins | Jaramillo et al. 2017, Bicudo et al. 2019, Bernal et al. 2019 |
| **W-Amazon** | 1 | 1 | 0 | 1 | 1 | 0 | Rise of Bolivian Orocline captured seasonally some headwater of La Plata drainages by the Upper Madeira River (Western-Amazon) | Tagliacollo et al., 2015 |
| **E-Amazon** | 1 | 0 | 1 | 1 | 0 | 0 | | |
| **Pacific Coast** | 1 | 1 | 1 | 1 | 0 | 0 | | |
| **La Plata** | 0 | 1 | 0 | 0 | 1 | 1 | Rise of Serra do Mar: partial disconnection between La Plata drainage basins from southeastern basins | Ribeiro 2006, Cogne et al. 2012 |
| **Atlantic Coast** | 0 | 0 | 0 | 0 | 1 | 1 | | |
| **Late Miocene to Recent: 10 – 0 Ma** | | | | | | | Diversification in modern trans-continental Amazon | |
| **Guianas** | 1 | 0 | 1 | 0 | 0 | 0 | Rise of Vaupes Arch: disconnected Western Amazon and Guianas (with intermittent connection by Casiquiare channel) | Hoorn et al., 2010b |
| **W-Amazon** | 0 | 1 | 1 | 0 | 1 | 0 | Rise of Northern Andes / Breach Purus Arch: connected Western and Eastern-Amazon | Hoorn et al., 2010; Hoorn et al. 2017, Albert et al. 2018 |
| **E-Amazon** | 1 | 1 | 1 | 0 | 0 | 0 | Rise of Northern Andes: disconnected Pacific Coast from Guianas and Pacific Coast from Western-Amazon. | Albert et al. 2006 |
| **Pacific Coast** | 0 | 0 | 0 | 1 | 0 | 0 | | |
| **La Plata** | 0 | 1 | 0 | 0 | 1 | 1 | | |
| **Atlantic Coast** | 0 | 0 | 0 | 0 | 1 | 1 | | |

**Biogeographical Regionalization.** To better understand the evolutionary history of freshwater fishes South America was classified into six biogeographical regions (regions hereafter). The biogeographical regionalization process starts from 490 present-day drainage basins, at level 5, according to the Hydrobasin database (https://www.hydrosheds.org/page/hydrobasins). To classify these drainage basins into biogeographical regions we used the approach proposed by Kreft & Jetz (2013). Considering the uncertainty in the current distribution of freshwater fishes across the 490 basins, we used 100 potential presence/absence matrices (see main text) to calculate 100 pairwise "taxonomic" dissimilarity matrices (using G. G. Simpson's presence/absence similarity/dissimilarity index, Simpson 1943, Koleff et al. 2003). Likewise, we also used phylogenetic distance among species to calculate dissimilarity matrices among basins. As phylogenetic trees by Rabosky et al. (2018) present some taxonomic incongruences, we followed Hughes et al. (2018) and adjusted the relationships within Ostariophysi to (Characiformes, (Gymnotiformes, Siluriformes)). Then, we reduced the dimensionality of these dissimilarity matrices using NMDS. Visual inspection of the map of the first three NMDS axes (averages among 100 NMDS), as shown in the rasterized RGB plot of Figure S1, highlights conspicuous biogeographical regions, which are similar in large-scale pattern, whether using taxonomic or phylogenetic similarities.

Finally, from the taxonomic dissimilarity matrices, we used K-means to cluster the 490 basins into six biogeographical regions for freshwater fishes (view main text, Fig. 3). K-means clustering was also repeated 100 times, considering the 100 alternative species presence/absence matrices, but generated similar clusters of basins.

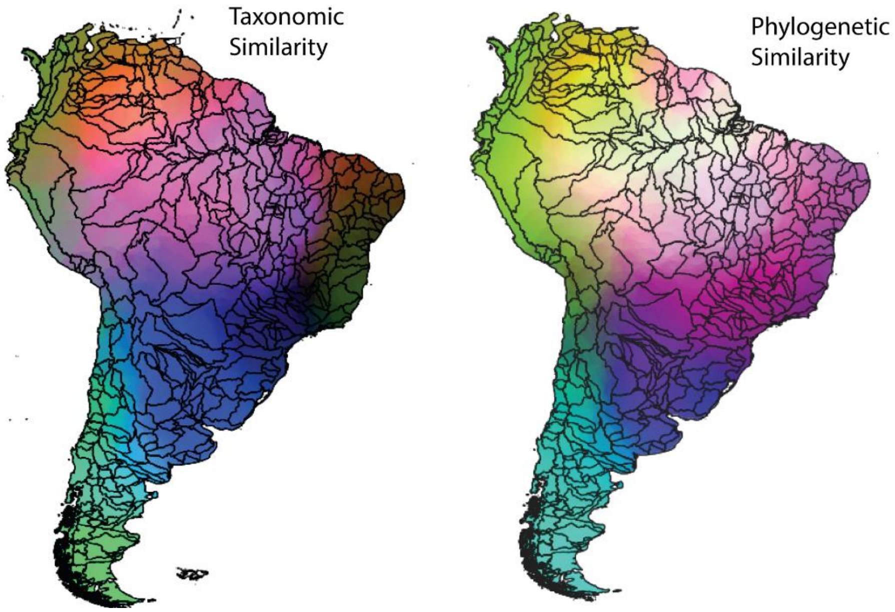

**Figure S1**. RGB interpolation of the three axes of NMDS based on 100 species presence/absence matrices (taxonomic similarity) and phylogenetic matrix (phylogenetic similarity).

**Table S2.** Proportion of dispersal events (%) among the six regions, considering all estimated dispersal events (17,244.2), which are an average of 20 simulations. Bold numbers represent the highest proportions of dispersal events from (lines) one region to (columns) another region. See also Table S3.

| Regions From/To | Guianas | W-Amazon | E-Amazon | Pacific Coast | La Plata | Atlantic Coast |
|---|---|---|---|---|---|---|
| **Guianas** | - | 4.1 | **6.4** | 4.1 | 0.4 | 0.5 |
| **W-Amazon** | 4.3 | - | 7.0 | 5.5 | **9.8** | 0.5 |
| **E-Amazon** | **6.8** | 5.7 | - | 4.8 | 1.7 | 0.5 |
| **Pacific Coast** | 4.1 | 4.8 | **5.6** | - | 0.5 | 0.5 |
| **La Plata** | 0.5 | 5.6 | 0.4 | 0.6 | - | **9.2** |
| **Atlantic Coast** | 0.5 | 0.4 | 0.5 | 0.6 | **4.0** | - |

**Table S3.** Average number of dispersal events (from 20 simulations) estimated by Biogeographical Stochastic Mapping analysis, using the BioGeoBEARS package. Bold numbers represent the highest values of dispersal from (lines) one region to (columns) another region.

| Regions From/To | Guianas | W-Amazon | E-Amazon | Pacific Coast | La Plata | Atlantic Coast | Total | % |
|---|---|---|---|---|---|---|---|---|
| **Guianas** | - | 714.6 | **1105.85** | 709 | 65.85 | 79.3 | **2674.6** | 15.5 |
| **W-Amazon** | 736.5 | - | **1214.65** | 942.3 | **1684.3** | 80.2 | **4657.95** | **27.0** |
| **E-Amazon** | **1180.8** | 990.5 | - | 835 | 295.3 | 80.25 | **3381.85** | **19.6** |
| **Pacific Coast** | 712.6 | 831.4 | 971.25 | - | 80.55 | 80.35 | **2676.15** | 15.5 |
| **La Plata** | 79.55 | 972.6 | 76.9 | 100.95 | - | **1585.95** | **2815.95** | 16.3 |
| **Atlantic Coast** | 94.55 | 71.05 | 83.05 | 98.05 | **690.95** | - | **1037.65** | 6.02 |
| | | | | | | **Total** | **17244.2** | **100.0** |

**Evolutionary Distinctiveness**. We estimated the uniqueness of a species using the evolutionary distinctiveness measure (ED; Isaac et al. 2007). To calculate ED, the total phylogenetic diversity (PD) of a clade was split equally among its members, which estimates how much branch length, on average, is unique to each species. The ED of a species is the sum of these values for all branches from which the species is descended, to the root of the phylogeny. We calculated the average of ED by summing the branch length of all species and dividing by the total of species in each basin.

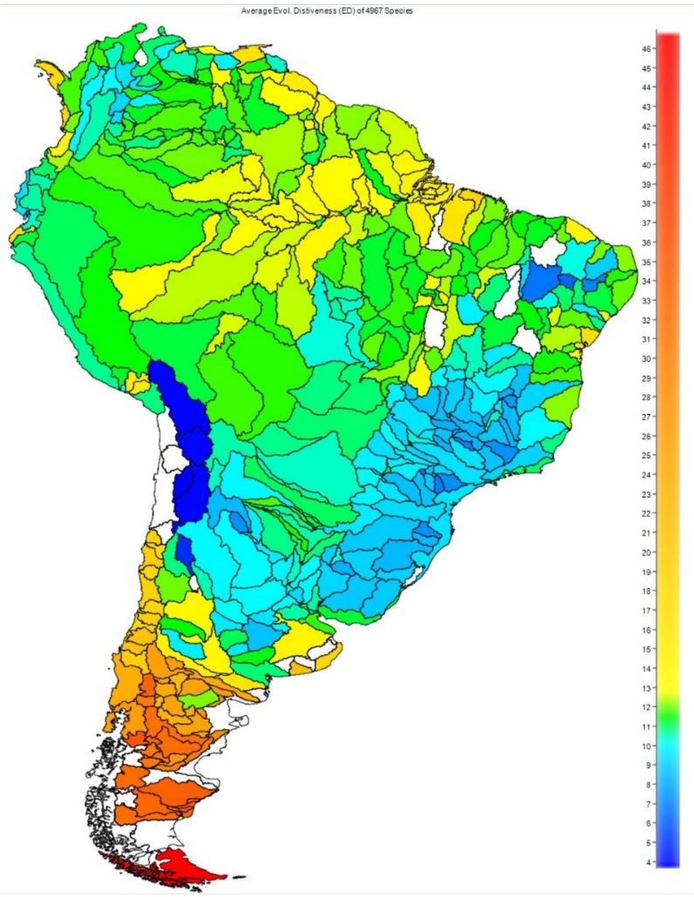

**Figure S2.** Average of Evolutionary Distinctiveness (ED) per basin. ED is measured as the distance along the evolutionary tree from one lineage to its nearest relative and shows the amount of unique evolutionary history a lineage represents. Data for 4,967 species in 490 basins. Basins in white are data deficient.

**Phylogenetic Endemism Analysis.** We conducted a phylogenetic endemism analysis (PEA; Rosauer et al. 2009) as a relative measure of endemism that apportions each unit of PD across the areas where it occurs. This measure identifies concentrations of taxa with spatially restricted PD (Faith et al., 2004; Cadotte & Davies, 2010). PEA uses the branch length and the geographic range of the extant descendants of each branch on a phylogeny to sum the proportion of the geographic range of each unit of PD found in an area (each of the 490 basins). This analysis considers the geographic range of each species (Crisp et al., 2001) and identifies concentrations of spatially restricted species. A phylogenetic dated tree for a set of 4,967 freshwater fish species was used (Rabosky et al. 2018) to calculate phylogenetic endemism as:

$$PE = \sum_{c \in C} \frac{L_c}{R_c},$$

where *C* is a phylogenetic clade spanned by minimal branches to link all taxa within one basin, and *c* is any branch between two nodes within *C*; $L_c$ is the length of branch *c*; and $R_c$ is the range occupied by branch *c*, defined in numbers of grid cells (Rosauer et al., 2009).

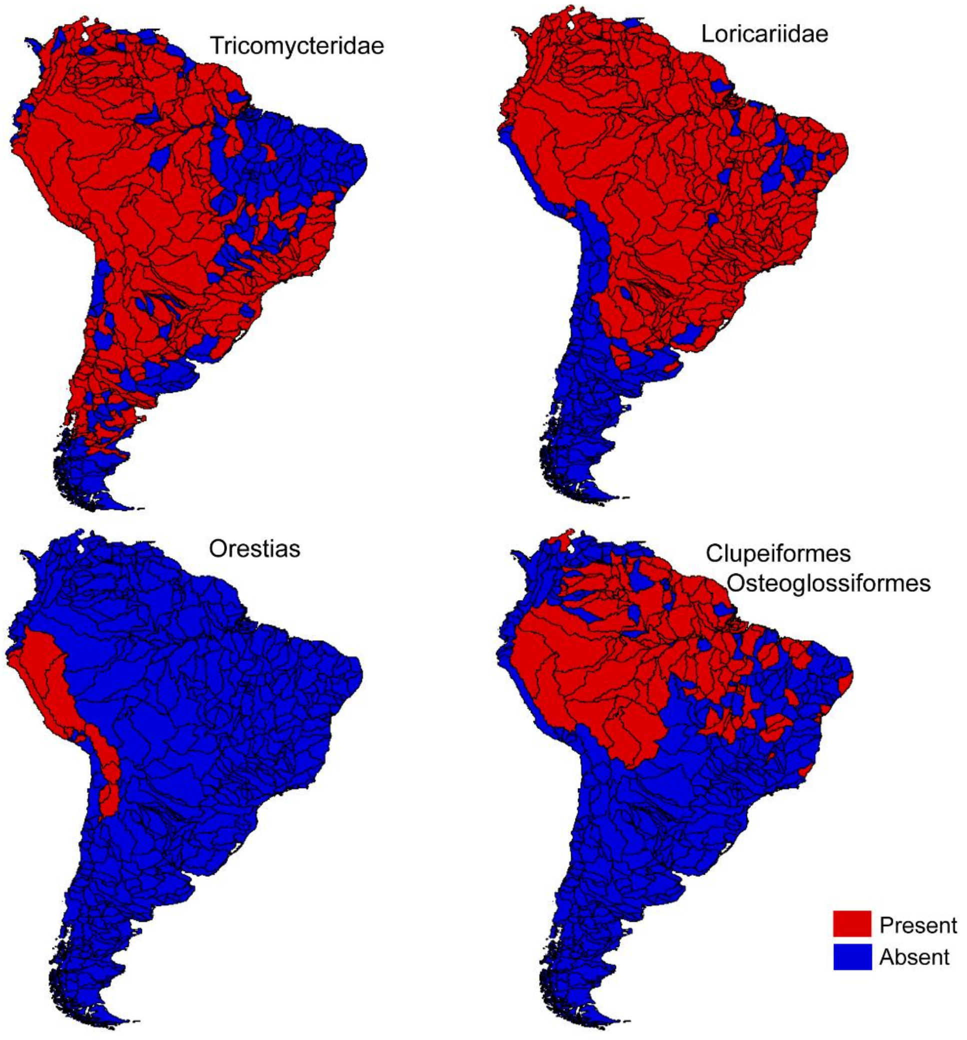

**Figure S3.** Current distribution (presence and absence) of the clades that showed rapid shifts in diversification rates.

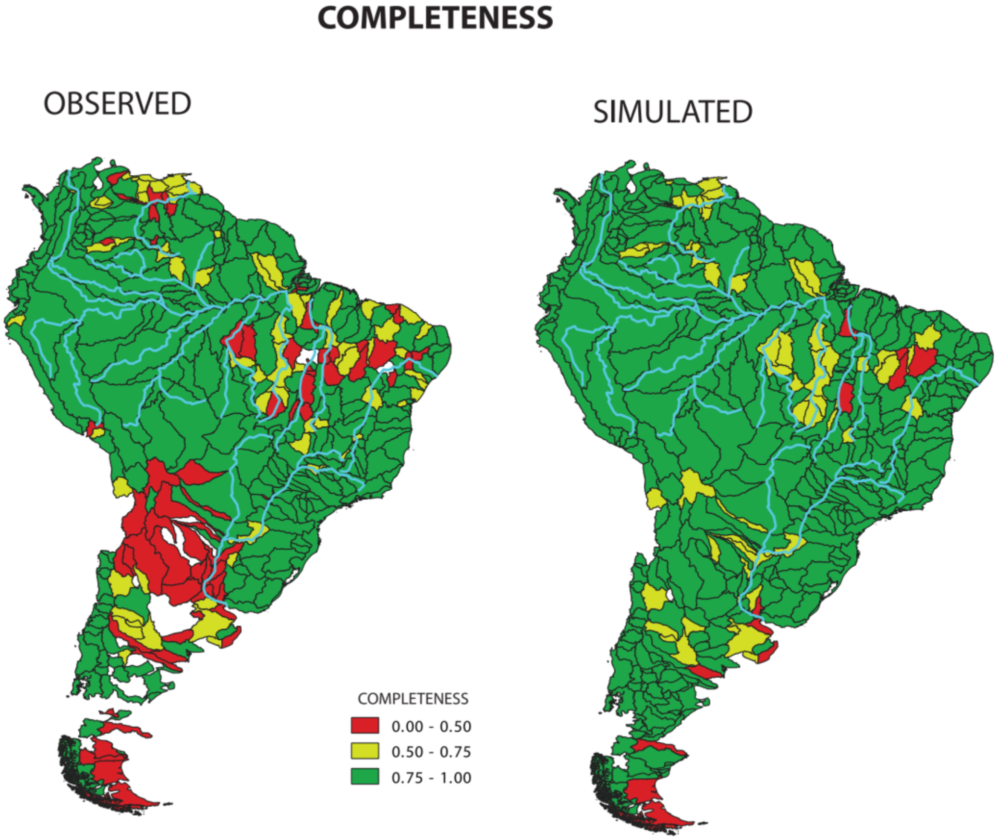

**Figure S4**. Delimitation of 490 drainage basins of level 5 (following the HydroSHEDS database) and sampling completeness at each basin. On the left, green indicates drainage basins with good sampling completeness, red and yellow basins indicate poor sampling completeness, and white shows basins without sampling records. In standard statistical analysis, white, red, and yellow basins would have to be excluded from the study because of poor sampling completeness. On the right, the map represents sampling completeness values after one simulation, in which species records are randomly sampled among level 5 basins that belong to the same level 4 basin (see main text). As the figure shows, the completeness values of several basins increased substantially (shifting from white, red, or yellow to yellow or green). Yet, even with simulation, several basins remained poorly sampled (red or white) and were excluded from analysis.

**Appendix 1:**

The area coding at the tips of the phylogenies depicts current distributions, while at internodes the coding represents the most likely ancestral area distribution as inferred under the Dispersal-Extinction-Cladogenesis (DEC) model implemented in BioGeoBEARS. Region abbreviations: A = Guianas, B = Western Amazon, C = Eastern Amazon, D = Pacific Coast, E = La Plata, and F = Atlantic Coast.
